# Selection in males purges the mutation load on female fitness

**DOI:** 10.1101/2020.07.20.213132

**Authors:** Karl Grieshop, Paul L. Maurizio, Göran Arnqvist, David Berger

**Affiliations:** Department of Ecology and Genetics, Animal Ecology, Uppsala University, Uppsala, Sweden; Department of Ecology and Evolutionary Biology, University of Toronto, Toronto, Canada; Department of Molecular Biosciences, The Wenner-Gren Institute, Stockholm University, Stockholm, Sweden; Section of Genetic Medicine, Department of Medicine, University of Chicago, Chicago, Illinois, United States

**Keywords:** Diallel cross, Fitness, Good genes, Heterosis, Mutation load, Sexual selection

## Abstract

Theory predicts that the ability of selection and recombination to purge mutation load is enhanced if selection against deleterious genetic variants operates more strongly in males than females. However, direct empirical support for this tenet is limited, in part because traditional quantitative genetic approaches allow dominance and intermediate-frequency polymorphisms to obscure the effects of the many rare and partially recessive deleterious alleles that make up the main part of a population’s mutation load. Here, we exposed the partially recessive genetic load of a population of *Callosobruchus maculatus* seed beetles via successive generations of inbreeding, and quantified its effects by measuring heterosis – the increase in fitness experienced when masking the effects of deleterious alleles by heterozygosity – in a fully factorial sex-specific diallel cross among 16 inbred strains. Competitive lifetime reproductive success (i.e. fitness) was measured in male and female outcrossed F_1_s as well as inbred parental ‘selfs’, and we estimated the 4×4 male-female inbred-outbred genetic covariance matrix for fitness using Bayesian Markov chain Monte Carlo simulations of a custom-made general linear mixed effects model. We found that heterosis estimated independently in males and females was highly genetically correlated among strains, and that heterosis was strongly negatively genetically correlated to outbred male, but not female, fitness. This suggests that genetic variation for fitness in males, but not in females, reflects the amount of (partially) recessive deleterious alleles segregating at mutation-selection balance in this population. The population’s mutation load therefore has greater potential to be purged via selection in males. These findings contribute to our understanding of the prevalence of sexual reproduction in nature and the maintenance of genetic variation in fitness-related traits.

**Impact statement:** Why do the large majority of eukaryotic species reproduce sexually if it means that females must spend half of their reproductive effort producing males, while males contribute few or no resources to offspring production themselves? In principle, a lineage of a mutant asexual female that simply clones herself into daughters would grow at twice the rate of her sexual competitors (all else equal). What prevents this from being the predominant mode of reproduction throughout eukaryotes? One hypothesis regards the role of males in facilitating the purging of deleterious mutations from the population’s genome since very strong selection in males, unlike selection in females, can occur in many species without reductions in population offspring numbers. Due to the inherent difficulties of isolating this source of standing genetic variation for fitness, empirical evidence for this theory is mixed and limited to indirect evidence from manipulative experiments and experimental evolution studies. Here we demonstrate that recessive deleterious alleles in a population of the seed beetle, *Callosobruchus maculatus*, are selected against strongly in males but not females. Using a fully factorial diallel cross among 16 inbred strains, we measured the degree to which fitness in the outbred offspring of those crosses improved relative to their inbred parents. This measure is known as heterosis and offers an estimate of the relative amount of partially recessive deleterious alleles carried by a genetic strain. We then analyzed the relationship between the strains’ heterosis values and their additive genetic breeding values for fitness measured in males and females, revealing the extent to which segregating (partially recessive) deleterious alleles are selected against in males and females. We found that a strain’s heterosis value was strongly genetically correlated with its additive genetic breeding value for male fitness, but not female fitness. This suggests that mutations with deleterious effects on population growth rate due to their effects on females can be selected against (i.e. purged) more efficiently via their male siblings. This process would offer a benefit to sexual reproduction that may partly compensate for its costs, and therefore yields insight to the prevalence of sex in nature.

## Introduction

Sexual reproduction is paradoxically prevalent when considering the cost of producing male offspring, which contribute little to offspring production themselves (Maynard Smith 1971, 1978, Lehtonen et al. 2012, Gibson et al. 2017). Counterintuitively, this same male feature may offer long-term benefits to lineages producing sons, as deleterious alleles can be purged via strong selection in males without appreciable reductions to a lineage’s offspring production (Manning 1984, Kodric-Brown and Brown 1987, Agrawal 2001, Siller 2001, Whitlock 2002, Lorch et al. 2003, Whitlock and Agrawal 2009). For this process to advantage sexual lineages over mutant asexual competitors, and thereby account for the maintenance and prevalence of sex in eukaryotes, purifying selection against mutations with deleterious effects on female fecundity, and hence population offspring production, must be stronger in males than in females (Agrawal 2001, Siller 2001, Whitlock and Agrawal 2009). Selection is likely stronger in males than females in many systems (Wade and Arnold 1980, Whitlock and Agrawal 2009) owing to sexual selection operating more strongly in males (Wade 1979, Andersson 1994, Janicke et al. 2016), which is ultimately due to sex-differences in gamete investment (i.e. anisogamy; Parker et al. 1972, Schärer et. al. 2012).

Empirical support for male-enhanced purging of the genetic load on females comes mostly from studies of induced, accumulated or known mutations (e.g. Radwan 2004, Sharp & Agrawal 2008, 2013, Hollis et al. 2009, Grieshop et al. 2016) or experimental evolution (e.g. Firman and Simmons 2010, 2012, Lumley et al. 2015, Dugand et al. 2018, 2019a, Yun et al. 2018, Buzatto and Clark 2020, Kyogoku and Sota 2020; reviewed in Cally et al. 2019), but detecting this process in a snapshot of the standing genetic variation has proven difficult (Chenoweth et al. 2015). This difficulty may owe to interference between signals stemming from the ‘mutation load’ and the ‘segregation load’ (Crow 1958, Whitlock and Davis 2011, Charlesworth and Charlesworth 2010). The former is attributed to rare, often partially recessive, deleterious alleles in mutation-selection balance (Haldane 1927, Lande 1975, Lynch et al. 1999, Zhang X.S. et al. 2004) and the latter is attributed to (net) heterozygote advantage, including genetic tradeoffs where heterozygotes are the most fit on average due to alleles having conditionally deleterious/beneficial effects on fitness (Rose 1982, Connallon and Clark 2012). The fact that mutation-selection balance alone tends to be insufficient to account for all of the observed genetic variance in fitness and life history traits (Houle et al. 1994, Charlesworth and Hughes 2000, Barton and Keightley 2002, Mitchell-Olds et al. 2007, Charlesworth 2015, Sharp and Agrawal 2018) suggests that the segregation load comprises a considerable fraction of a population’s fitness variance. Thus, while sex-specific estimates of variance in fitness would indicate the relative strength of selection in males versus females (Crow 1958, Charlesworth and Hughes 2000, Cox and Calsbeek 2009, Janicke et al. 2016, Singh and Punzalan 2018), simply comparing fitness variance between the sexes would confound the effects of rare unconditionally deleterious alleles with those that impose conditional fitness effects. This is particularly problematic for comparing the strength of purifying selection between the sexes in light of segregating sexually antagonistic alleles, whose fitness effects are conditional upon sex (Chippindale et al. 2001, Bonduriansky and Chenoweth 2009, van Doorn 2009, Connallon et al. 2010, Connallon and Clark 2012). This is because strongly selected male-benefit/female-detriment alleles that impose detriments to female offspring production will be overrepresented in the standing genetic variation relative to alleles with detrimental effects in both sexes that are maintained at mutation-selection balance (Pischedda and Chippindale 2006, Long et al. 2012, Berger et al. 2016a). Indeed, this phenomenon is predicted to obfuscate experimental detection of the long-term benefits of male-enhanced purging of mutation load (Whitlock and Agrawal 2009, p. 576).

To address this issue, we experimentally uncovered sex-specific fitness effects of the rare, partially recessive deleterious alleles that comprise the mutation load in a population of the seed beetle, *Callosobruchus maculatus*, by increasing genome-wide homozygosity in 16 genetic strains and then analyzing heterosis for fitness (competitive lifetime reproductive success). Here, heterosis of a genotype is the increase in fitness in outbred progeny of crosses involving that genotype relative to its inbred/homozygous state (Charlesworth and Willis 2009). Whereas fitness variance (Charlesworth and Hughes 2000, Kelly and Willis 2001, Barton and Keightley 2002, Kelly 2003, Mitchell-Olds et al. 2007, Charlesworth 2015, Sharp and Agrawal 2018) and inbreeding depression (Charlesworth and Charlesworth 1987, 1999, Charlesworth et al. 2007, Dugand et al. 2019b) can both be attributable to the mutation load and the segregation load, heterosis in crosses among inbred strains of a given population should be disproportionately attributable to the masking of rare, partially recessive deleterious alleles by heterozygosity (Charlesworth and Willis 2009). Our estimates of strain-specific heterosis – the difference between a strain’s outbred and inbred fitness – should therefore primarily reflect each strain’s relative share of the population’s mutation load on female offspring production. While the segregation load may in theory also contribute to heterosis (Charlesworth and Willis 2009), it is unlikely to play a role in the present study population. First, the correlation between male and female heterosis is near unity (see *Results*), indicating that this population’s most relevant source of segregation load, its sexually antagonistic genetic variation (Berger et al. 2016, 2016a, Grieshop and Arnqvist 2018), contributes little or nothing to variation in heterosis among strains. Second, fitness in the inbred state exhibits a substantially reduced mean and increased variance relative to the outbred state (see *Results*), which is consistent with fitness being underlain by rare, partially recessive deleterious alleles (Robertson 1952, Houle et al. 1997, Kelly 1999, Charlesworth and Hughes 2000, Kelly and Tourtellot 2006).

We therefore reason that an approximation of the strength of purifying selection against mutation load alleles in each sex is given by quantifying the relationship between strains’ male and female additive genetic breeding values for relative fitness and their genetic values for heterosis (i.e. a measure that should reveal the relative amount of mutation load alleles carried by each strain). If selection in males acts to purge mutation load on female fecundity, there should be a negative genetic covariance between outbred male fitness and female heterosis (see *Methods*). In other words, strains with a greater relative share of the population’s mutation load should have lower male fitness. Similarly, if selection in females acts to purge mutation load, there should be a negative genetic covariance between outbred female fitness and heterosis. Thus, we predicted a negative relationship between heterosis and outbred fitness among strains, and that this relationship would be stronger in males than females if selection in males enhances the purging of mutation load.

## Methods

### Study population and inbred strains

*Callosobruchus maculatus* (Coleoptera: Bruchidae) is a pest of leguminous crops that has colonized most of the tropical and subtropical regions of the world (Southgate 1979). Laboratory conditions and fitness assays closely resemble the grain storage facilities and crop fields to which they are adapted. Females lay eggs on the surface of dry beans and hatched larvae bore into the beans, where they complete their life cycle, emerging from the beans as reproductively mature adults (Southgate 1979). This species is facultatively aphagous (requiring neither food nor water to reproduce successfully), exhibits a generation time of ~3 weeks (Southgate 1979), and exhibits a polyandrous mating system (Miyatake and Matsumura 2004, Katvala et al. 2008).

The origin of this population’s isofemale lines and inbred lines have been described thoroughly by Berger et al. (2014), Grieshop et al. (2017), and Grieshop and Arnqvist (2018). Briefly, 41 isofemale lines were constructed from a wild population of *C. maculatus* that was isolated from *Vigna unguiculata* seed pods collected at a small-scale agricultural field close to Lomé, Togo (06°10’N 01°13’E) during October and November 2010 (Berger et al. 2014; see Supporting Information Fig. S1A). Each isofemale line stemmed from a single virgin male/female pair whose offspring were expanded into small sub-populations (Fig. S1A). These 41 isofemale lines have an inbreeding coefficient of 0.25 (Falconer and Mackay 1996), and were cultured for 12 generations prior to the fitness assays of Berger et al. (2014). From January 2013 to January 2014, 20 replicate lineages of each isofemale line (totaling >800 lineages) were subjected to single-pair full-sibling inbreeding (i.e. “close” inbreeding; Falconer and Mackay 1996) for 10 consecutive generations or until extinction (Grieshop et al. 2017; Fig. S1A). For the 16 inbred strains chosen for the present diallel cross, this was followed by one generation of expansion into small populations, comprising a total of 12 generations of full-sibling mating (1 full-sibling isofemale line expansion + 10 generations of close inbreeding + 1 full-sibling inbred line expansion), which corresponds to an inbreeding coefficient of 0.926 (Falconer and Mackay 1996). During the inbreeding regime, inbred lineages stemming from isofemale lines that were enriched for male-benefit/female-detriment sexually antagonistic genetic variation tended to go extinct prior to completing the full inbreeding program (Grieshop et al. 2017), making it impossible to retrieve a representative inbred line from four of the most male-benefit/female-detriment isofemale lines. The present 16 inbred strains were chosen with the aim of countering that biased representation (Fig. S2).

### Sex-specific fitness assay

The present study used data from a fully factorial diallel cross (Lynch and Walsh 1998) among the 16 inbred strains, where sex-specific competitive lifetime reproductive success (hereafter: fitness) was measured in F_1_ males and females separately. The partitioning of genetic variance in fitness is reported in detail by Grieshop and Arnqvist (2018), and the aspects that bear relevance to the present study are given below and in the *Discussion*. The experiment was conducted in two replicate ‘blocks’ for a total of 3278 individual fitness estimates performed in 237/240 possible outbred crosses and all 16 parental selfs (Fig. S1B). Male and female fitness, as well as inbred and outbred fitness, were measured independently (Fig. S1B,C), and all were approximately normally distributed. There are 1616 outbred male (*o_M_*), 1450 outbred female (*o_F_*), 115 inbred male (*i_M_*), and 97 inbred female (*i_F_*) individual fitness estimates. Each observation of the fitness assay consisted of a 90 mm ø petri dish containing ca. 100 *V. unguiculata* seeds, a focal individual from a given outbred cross or inbred self, a sterilized same-sex competitor from the outbred base population, and two opposite-sex individuals from the base population (Fig. S1C). Same-sex competitors were sterilized with a 100 Gy dose of ionizing radiation from a Cs^137^ source (Fig. S1C), which does not notably reduce lifespan or reproductive competitiveness in either sex, but does cause zygote lethality, accrediting all emerging offspring to focal individuals (Grieshop et al. 2016). Thus, counts of F_2_ offspring emerging in the petri dishes are integrative measures of focal individuals’ fitness (Fig. S1C), the differences among them being attributable to focal F_1_ individuals’ pre- and post-copulatory reproductive success and their offspring’s larval viability. Our fitness assays therefore enable, but are not limited to, the following mechanisms of selection in adults: mate searching and oviposition site selection, mating success, mating resistance, sexual conflict over remating rate, sperm competition, cryptic female choice, fertility, fecundity, lifespan and offspring egg-adult survival (discussed further by Grieshop et al. 2016, Grieshop and Arnqvist 2018). Despite lacking many of the elements of natural selection that might apply to these beetles in nature (as would any laboratory fitness assay), this method has been effective in revealing sex-specific genetic variance in fitness (Berger et al. 2014, 2016a,b, Grieshop et al. 2016, Martinossi-Allibert et al. 2019a), perhaps owing to its relatively greater resemblance to the beetles’ natural ecology compared with other model systems and/or the inherent three-dimensional physical complexity provided by the beans, which may play an important role in achieving balance between sexually concordant and sexually antagonistic mating interactions in the laboratory (Singh et al. 2017, Yun et al. 2017).

### Modelling the genetic covariance matrix

We modelled the male-female-inbred-outbred additive genetic (co)variance matrix for fitness, **H**:

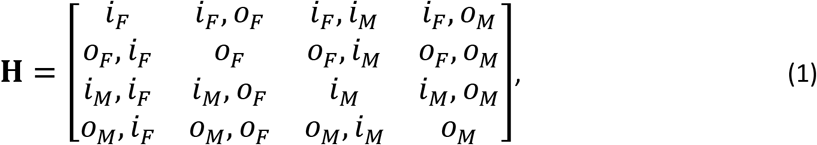

where genetic variances were estimated for parameters listed along the diagonal, and genetic covariances were estimated for pairs of parameters listed in the off-diagonal elements. While this resembles a cross-sex cross-trait **G** matrix that would typically be modeled with the aim of assessing the sex-specific genetic architecture of multiple traits and whether it constrains or enables (sex-specific) adaptation (Lande 1980, Gosden and Chenoweth 2014, Ingleby et al. 2014, McGlothlin et al. 2019, Sztepanacz and Houle 2019, Cheng and Houle 2020), our **H** matrix is distinct in two important ways: (1) it is a cross-sex cross-*state* (rather than cross-trait) (co)variance matrix, which models inbred/homozygous effects versus outbred/heterozygous effects for the same “trait”, and (2) that “trait” is fitness.

The **H** matrix was modeled in a general linear mixed-effects model (GLMM) using Bayesian Markov chain Monte Carlo (MCMC) simulations in the ‘MCMCglmm’ package (v. 2.25; Hadfield 2010) for R (v. 3.6.0; R Core Team 2019). To attain proper estimates of additive genetic variance, two additional random effects (and corresponding variance components) were included to estimate symmetrical epistasis (*v*) and sex-specific symmetrical epistasis (*v*×*S*) (i.e. (sex-specific) strain-strain interaction variance among outcrossed families only), as these effects are known to be present in these data (Grieshop and Arnqvist 2018). Fixed factors in this model were sex (*S*, male or female), inbred (*I*, inbred self or outbred cross), block (*B*, first or second replicate of the full diallel cross), and the interactions *S*×*I* and *B*×*I*, making the full GLMM:

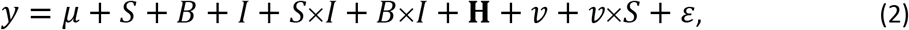

where *y* is relative fitness, *μ* is the intercept, and *ε* is the residual/unexplained error, normally distributed as *ε* ~ *N*(0, *σ*^2^) with variance *σ*^2^. We enabled (co)variance estimation to differ among elements of the **H** matrix, and used minimally informative parameter-expanded priors (Hadfield 2012). The model was run for 2,000,000 iterations after a burn-in of 200,000, with a thinning interval of 2000, which provided 1000 uncorrelated posterior estimates of each sex-/strain-specific effect to be stored and used for resampling the relationships described below. We used the Gelman-Rubin criterion to ensure model convergence (Gelman and Rubin 1992; Fig. S3). Posteriors were unimodally distributed and their trend stable over the duration of the simulations after the burn-in period. The model that was fit for the purpose of estimating the **H** matrix and resampling the stored posteriors (see below) was fit to relative fitness, i.e. fitness standardized by the sex-specific outbred mean, whereas the model that was fit for the purpose of plotting results was fit to untransformed/raw data.

We estimated heterosis, e.g. in females, as the difference between a strain’s outbred, *o_F_*, and inbred, *i_F_*, fitness. As explained in the *Introduction* and in more detail below, the genetic variance in female heterosis, *V*(*o_F_* − *i_F_*), and its genetic covariance with male outbred fitness, *COV*(*o_M_*, *o_F_* − *i_F_*), are of central interest to our study, but are not explicitly modeled in our **H** matrix. Relationships involving heterosis were therefore assessed by resampling the sex-/state-specific breeding values from the 1000 stored posteriors. This approach incorporates the uncertainty around those breeding values into the estimated relationships and their credibility intervals, hence enabling the quantification of statistical significance in a way that accounts for breeding values being otherwise anticonservative when used to assess relationships that are not accounted for by the model (Postma 2006, Hadfield et al. 2010).

### Estimating genetic relationships between outbred fitness and heterosis

Assuming that heterosis is predominantly due to rare (partially recessive) deleterious alleles (see *Introduction* and *Discussion*), we develop two complementary measures of how such mutation load alleles affect outbred fitness. Our first measure approximates the strength of selection on partially recessive deleterious alleles 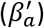. In males, for example, 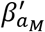 is given by the genetic covariance between outbred relative fitness and female heterosis, divided by the genetic standard deviation in heterosis: 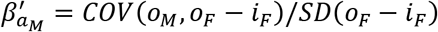. We note that our measure of selection is an additive genetic version (hence the subscripted ‘*a*’) of a univariate standardized phenotypic selection gradient (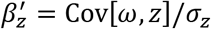, Lande & Arnold 1983, p. 1219), or “selection intensity” (Crow 1958, Falconer and MacKay 1996), used to provide comparable estimates of the strength of phenotypic selection across different traits or populations (Matsumura et al. 2012, Lynch & Walsh 2018). Our usage of this formulation differs from its original application in that our ‘phenotype’, heterosis, is not a feature of individuals but rather of genetic strains, and hence, our estimate of selection intensity is based on genetic (co)variances as advocated by Rausher (1992). This estimate – being standardized by *SD*(*o_F_* − *i_F_*) – has an advantage to other measures of selection as it is independent of the arbitrary magnitude of heterosis in our population, which is a direct consequence of the number of generations of inbreeding that we applied in our experiment. Thus, 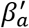 gives the genetic change in outbred relative fitness associated with one genetic standard deviation change in heterosis, which reflects the amount of mutation load alleles in the present population. Further, the comparison of this standardized measure of selection in males 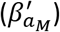 versus females (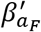, defined below) estimates the sex difference in selection against these mutation load alleles. Retrieving unbiased estimates of 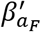 is problematic, however, due to measurement error in female outbred fitness also featuring in the heterosis term of both the numerator and denominator, which could drive spurious correlations and false positive discoveries (Postma 2011, Berger and Postma 2014). To address this, we explored whether male estimates of heterosis (*o_M_* − *i_M_*) may be so highly genetically correlated to the female estimates (*o_F_* − *i_F_*) that they effectively convey the same information, enabling 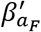 to be estimated using male heterosis. As male and female heterosis were, indeed, highly correlated (see *Results*), we ultimately estimated selection in males and females as:

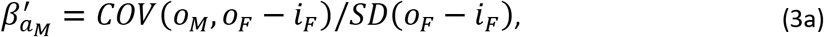

and

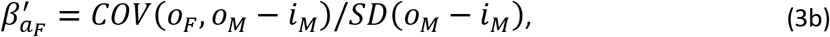

respectively.

The genetic correlation between male and female heterosis, *r_OM−iM,OF−iF_* (used to assess the validity of 3a and 3b), is also informative regarding the degree to which the deleterious effects of the genetic variation underlying heterosis in our population are conditional upon sex or not, where *r_OM−iM,OF−iF_* = 1 would indicate that heterosis is attributable to alleles whose effects are completely unconditional on sex and correlations below unity would indicate some sex-specificity to heterosis. Thus, the following genetic correlation was resampled 1000 times from the stored posteriors:

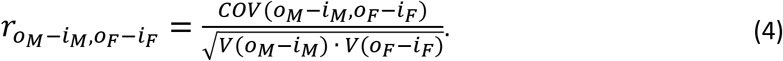

However, even though *o_M_* − *i_M_* and *o_F_* − *i_F_* were highly genetically correlated (see *Results*), 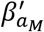 and 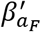 are not directly comparable. Thus, the assessment of whether 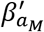 and 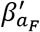 were significantly different, as well as the magnitude of their fold difference, was assessed using sex-averaged heterosis, i.e. 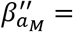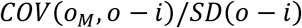 and 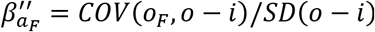, where 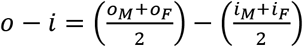. Despite the potential bias owing to shared measurement error (described above), 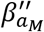 and 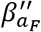 provided a qualitatively identical result to 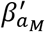 and 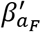, respectively (see Table S1), but enable a like-to-like comparison that is our least caveated estimate of the sex-difference in the efficacy of purifying selection against partially recessive deleterious alleles.

Our second estimate of how these rare partially recessive deleterious alleles affect fitness is given by the genetic correlation between outbred fitness and heterosis. The issue with shared measurement error in female outbred fitness and heterosis also applies to these correlations, and we thus took the same approach as described above when estimating the male and female correlations:

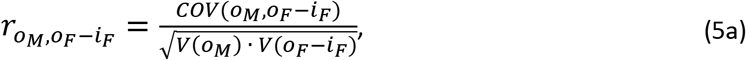

and

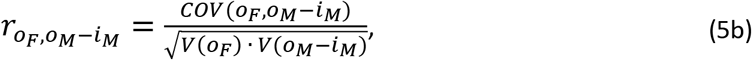

respectively. These correlations are merely different standardizations of the same numerator as the 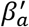 estimates (above), hence, their predictive frameworks (and reasoning therein) are similar to that given above, but with some important differences. Most importantly, they no longer reflect the strength of selection, *per se*, as they are standardized by fitness variance. Thus, these correlations represent the extent to which outbred fitness variance reflects mutation load alleles in each sex, where a correlation of −1 suggests that all fitness variance is solely due to rare and unconditionally deleterious alleles. These correlations are thus of particular interest for sexual selection theories of mate choice (Zahavi 1975, Hamilton & Zuk 1982, Grafen 1990, Andersson 1994) and “genic capture” in sexually selected traits (Rowe & Houle 1996, Tomkins et al. 2004), the latter based on the specific assumption that variance in sexually selected traits is maintained by polygenic deleterious mutation (see *Discussion*). Again, sex-averaged heterosis (see above) was used to assess whether male and female genetic correlations were significantly different from one another. Our main interpretations are therefore based on the estimated selection intensities (equation 3a & 3b) and genetic correlations (equation 5a and 5b), where sex differences in those were assessed using sex-averaged heterosis. All frequently used symbols are listed in Table A1.

Note that the selection intensities and correlations that were based on sex-averaged heterosis not only feature shared measurement error between outbred fitness and heterosis, but potentially also shared MCMC sampling error upon resampling these estimates from the posteriors. To avoid this issue, each of the 1000 resampled estimates of sex-averaged selection intensities and correlations drew their vector of sex-specific outbred fitness estimates (*o_M_* and *o_F_*) from different iterations than those used for sex-averaged heterosis (“*o* − *i*”). The point estimates of these selection intensities and genetic correlations are the posterior mode of the Bayesian posterior distribution based on 1000 resampled estimates, and these distributions were unimodal in all cases. The (95%) credibility intervals (CIs) around those point estimates are given by the highest posterior density (HPD) intervals. Two-tailed *P* values for these correlations and covariances were calculated as the proportion of times that those 1000 estimates fell on the opposite side of zero relative to the posterior mode (or overlapped the point estimate of the other sex in the case of assessing sex differences), multiplied by two. The plotted breeding values, heterosis estimates, and 95% confidence ellipses are for visual purposes only, and are based on the HPD means of the model fit of untransformed/raw fitness; they do not depict the uncertainty in those breeding values that was incorporated into the resampled estimates of CIs and *P* values. Because these HPD means are zero-centered in the ‘MCMCglmm’ output (even for models fit to untransformed data), they were rescaled to the more intuitive original scale before plotting, and the minimum heterosis value was set to zero.

The potential for non-genetic parental effects and sex-chromosome inheritance to explain our findings was thoroughly addressed. In short, we statistically removed these effects from our data and re-ran our analyses to confirm that our findings stand in the absence of those effects (Appendix A2). See *Data availability* regarding the R code for reproducing all analyses, procedures, and tables.

## Results

The genetic variance in fitness for inbred males, *V*(*i_M_*), was 1.85x that of inbred females, *V*(*i_F_*), and the genetic variance for outbred males, *V*(*o_M_*), was 1.24x that of outbred females, *V*(*o_F_*) (Table 1). Males also exhibited 3.37x and 5.58x the residual variance in fitness relative to females when inbred and outbred, respectively (Table 1). Genetic variance for inbred fitness was 10.92x and 7.35x that of outbred fitness in males and females, respectively (Table 1).

**Table 1.**
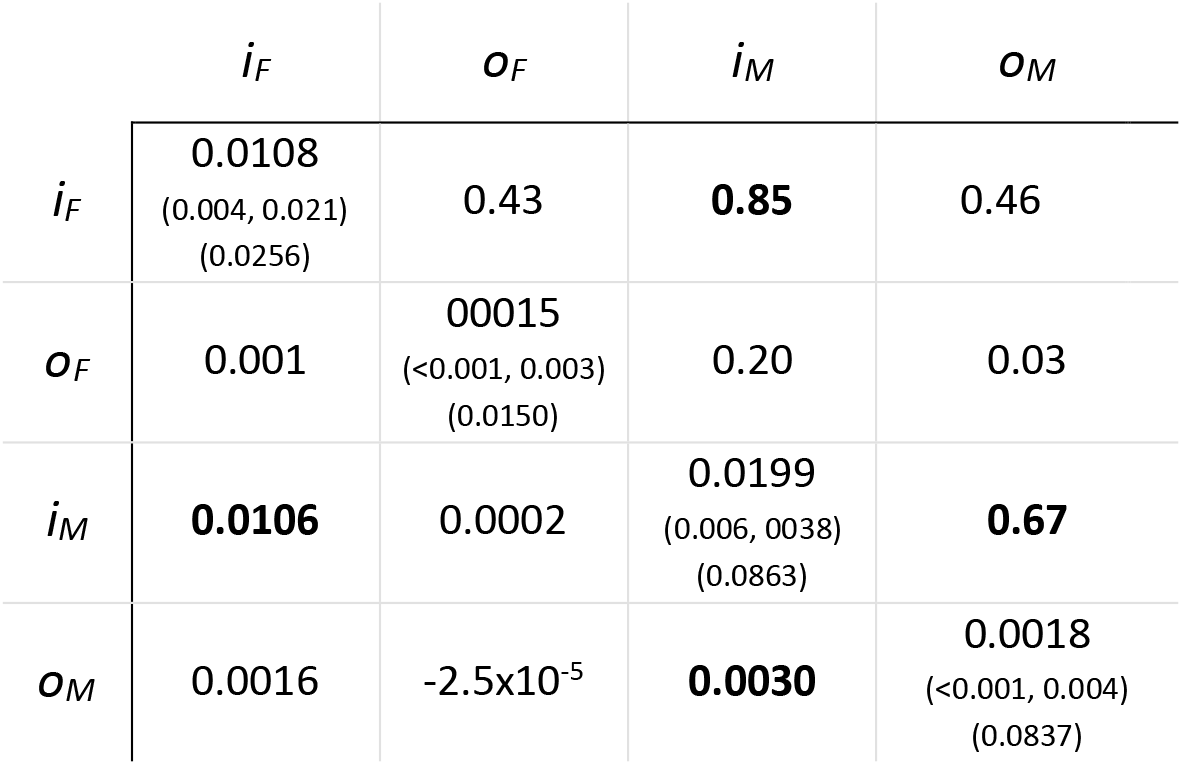
Estimated **H** matrix for relative fitness, displaying genetic variances (with credibility intervals (CIs) and residual variances) on the diagonal, their covariances in the lower triangle, and their Pearson’s correlation coefficients (*r*) in the upper triangle. Covariances and correlations with 95% credibility intervals excluding zero are bolded.

Whereas outbred fitness in males and females was genetically uncorrelated (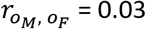, (95% CIs: −0.56, 0.54); Fig. 1A, Fig. S4), inbred fitness was highly genetically correlated (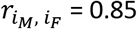 (0.38, 0.97); Fig. 1A, Fig. S5), suggesting that heterosis effects were large and unconditionally deleterious with respect to sex (estimated directly below). These correlations are explicitly estimated in the **H** matrix (Table 1). There was a large global improvement in mean fitness of outcrossed observations relative to the inbred/homozygous parental selfs (i.e. the fixed effect of *I*, mean reduction in relative fitness of inbreds versus outbreds = 0.294 (0.19, 0.41), *P*_MCMC_ < 0.001; Fig. 1A).

**Fig. 1.**
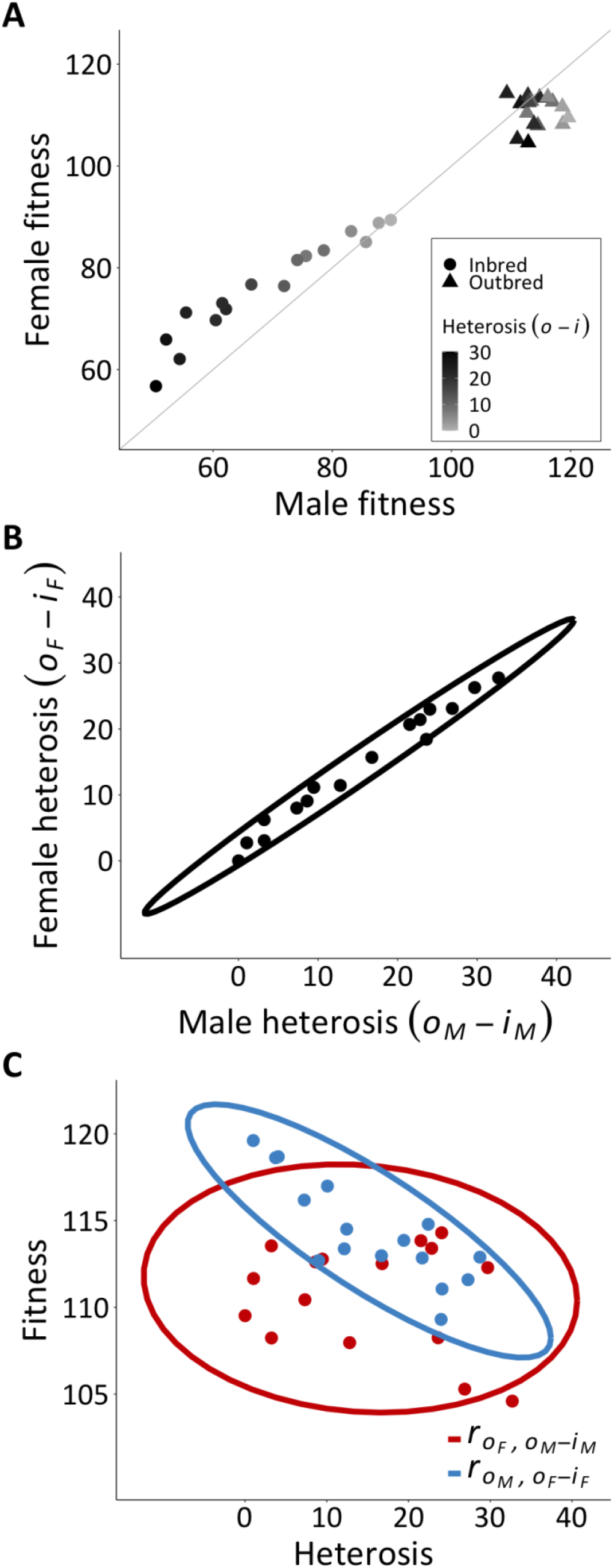
Breeding values from the MCMC model of untransformed (raw) fitness (for plotting purposes only). (A) A summary of the data and main result (y=x line for reference), showing each strain’s male (x-axis) and female (y-axis) fitness in the inbred (circles: 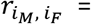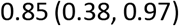) and outbred (triangles: 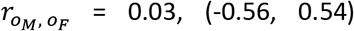) state, shaded by sex-averaged heterosis (i.e. ((*o_M_*-*i_M_*)+(*o_F_*-*i_F_*))/2). Variation in heterosis is clearly distributed along the population’s outbred male, but not female, breeding values. (B) Depiction of the genetic correlation for heterosis in male and female fitness across strains, showing that these sex-specific measures are conveying essentially the same information (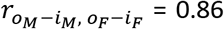, (0.66, 0.95), *P* < 0.001). (C) Depiction of main finding: the statistically significant resampled genetic correlation between outbred male fitness and female heterosis (blue: 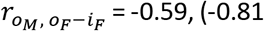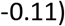, *P* = 0.008) would enable the mutation load on population mean fitness to be purged via selection in males. In contrast, the outbred female breeding values do not reflect this mutation load (red: 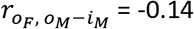, (−0.40, 0.28), *P* = 0.672). Ellipses are 95% confidence ellipses fit to the breeding values, and therefore do not depict the uncertainty that was included in the resampled estimates of statistical significance (see *Methods*). 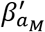 and 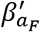 depicted in Fig. S6.

Heterosis was highly genetically correlated between males and females (resampled 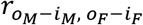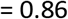, (0.66, 0.95), *P* < 0.001; Fig. 1B) with even narrower credibility intervals than the inbred intersexual genetic correlation, 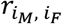 (see above), showing alignment between the sexes in the deleterious effects revealed by heterosis among strains. The point estimate of the strength of selection in males against these partially recessive deleterious effects on female fitness showed that one genetic standard deviation change in heterosis was associated with 1.25% reduction in outbred male fitness (resampled 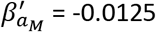, (−0.031, −0.003), *P* = 0.008; Fig. S6). By contrast, the corresponding estimate of selection in females was weak and undetectable (resampled 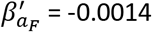, (−0.013, 0.009), *P* = 0.672; Fig. S6). Outbred male fitness was significantly and strongly negatively genetically correlated to female heterosis (resampled 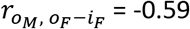, (−0.81, −0.11), *P* = 0.008; Fig. 1C), suggesting that a sizeable proportion of fitness variance in males is due to rare partially recessive deleterious alleles. Female fitness, by contrast, was not significantly genetically correlated to male heterosis (resampled 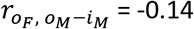, (−0.40, 0.28), *P* = 0.672; Fig. 1C). Estimates of selection and genetic correlations that were based on sex-averaged heterosis yielded qualitatively identical results (Table S1). Using those directly comparable estimates based on sex-averaged heterosis revealed that selection against partially recessive deleterious alleles was 3.7x stronger in males than females, yet estimates of selection (proportion of 1000 estimates of 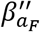 > 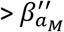 times two: *P* = 0.104) and genetic correlations (*P* = 0.12) were not significantly different between the sexes using two-sided hypothesis testing.

## Discussion

Our findings suggest that the mutation load of our population is more effectively purged via selection in males than in females (Fig. 1C). With heterosis being so highly sexually concordant (Fig. 1B), we were able to circumvent the potential bias caused by shared measurement error between fitness and heterosis by assessing the relationship between the outbred breeding values in one sex and heterosis in the other (see *Methods*, equations 3a,b and 5a,b). Moreover, relationships between male fitness and female heterosis (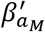 and 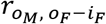; see equations 3a and 5a) are much more central to the question of whether selection via males can purge a population’s mutation load, since population productivity in most taxa is limited by female offspring production. That the female equivalents of this assessment (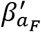 and 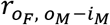; see equations 3b and 5b) were found to be indistinguishable from zero suggests that genetic variation in fitness among females of this population does not evidently reflect the partially recessive deleterious mutations that they carry in their genomes. These results remained essentially unchanged when analyses were performed on estimates of heterosis averaged across the sexes (Table S1).

We used heterosis as a measure of the relative share of the population’s mutation load that is captured within each of our strains. Heterosis among inbred strains of a population should predominantly owe to the same type of genetic variation that is often expected to constitute a population’s mutation load – rare, partially recessive deleterious alleles (Haldane 1927, Lande 1975, Houle et al. 1997, Lynch et al. 1999, Zhang X.S. et al. 2004, Charlesworth and Willis 2009). This is particularly likely in the case of our study population, as its genetic variance in fitness is characteristic of that underlain by rare, partially recessive deleterious alleles: fitness in the inbred state exhibits a substantially lower mean and greater variance relative to the outbred state (Robertson 1952; Houle et al. 1997, Kelly 1999; Charlesworth and Hughes 2000, Kelly and Tourtellot 2006; Table 1, Fig. 1A). Further, while this synthetic diallel population (Grieshop and Arnqvist 2018) as well as its wild-caught origins (Berger et al. 2014, 2016a) bear the hallmarks of fitness variance maintained by sexually antagonistic balancing selection, this sexually antagonistic genetic variation apparently plays little or no role in determining the magnitude of heterosis in crosses among inbred strains, as the magnitude of heterosis experienced on average among strains is nearly identical between the sexes (Fig. 1B). In accordance, a previous estimate of dominance variance in this population, which is based to a large extent on heterosis (Hayman 1954, Lynch and Walsh 1998, Lenarcic et al. 2012, Maurizio et al. 2018, Shorter et al. 2019), was likewise found to describe sexually concordant fitness effects (Grieshop and Arnqvist 2018). Reciprocally, the sex-reversed dominance effects (Kidwell et al. 1977, Fry 2010, Barson et al. 2015, Spencer and Priest 2016, Connallon and Chenoweth 2019) that were previously identified in this population (Grieshop and Arnqvist 2018) were detected via methods that are not based on heterosis. Thus, there is very little if any scope for this population’s genetic variance in heterosis to be attributable to factors other than rare, partially recessive deleterious alleles, and we are confident that the most relevant form of the ‘segregation load’ (see *Introduction*) that could possibly confound this interpretation – i.e. sexually antagonistic genetic variation – is absent from our estimates of sex-/strains-specific heterosis (further discussion in Appendix 3). The genetic covariance between those heterosis estimates and sex-specific outbred fitness therefore indicates how selection would act to purge the alleles that make up this population’s mutation load.

Our findings are pertinent to the longstanding question of why sexual reproduction is so prevalent in nature. Since, all else equal, the production of sons would halve the exponential growth rate of a sexual female’s lineage relative to an asexual competitor (Maynard Smith 1971, Lehtonen et al. 2012, Gibson et al. 2017), there must be some mechanism(s) that compensate for the two-fold cost of sex. One explanation is that the efficacy of selection against the mutation load on a population’s offspring production is greater in males relative to females, which would allow that load to be purged without the demographic costs that would ensue given that same strength of selection acting on the population’s females (Manning 1984, Kodric-Brown and Brown 1987, Agrawal 2001, Siller 2001, Whitlock 2002, Lorch et al. 2003, Whitlock and Agrawal 2009). Our most relevant estimate of the relative extent to which our population’s mutation load is purged via selection in males versus females is the fold difference between 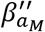 and 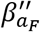. Although these estimates were not statistically significantly different from one another, the male estimate was highly significant while the female estimate clearly was not (as was the case for 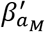 and 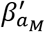; Table S1). As a rough guide, selection against rare, partially recessive deleterious mutations was estimated to be 3.7x greater in males than in females, in accordance with previous, more general, estimates of sex-specific selection in this species: ~3x stronger in males for selection against induced mutations (Grieshop et al. 2016) and 2-4x stronger for males in outbred populations (Fritzsche and Arnqvist 2013, Martinossi‐Allibert et al. 2020). We note that the fold difference of male-bias in selection that is needed in order for the production of males to yield a net benefit to females/populations does, however, depend on the genome-wide deleterious mutation rate, as well as other genetic, demographic and ecological factors (Agrawal 2001, Siller 2001, Agrawal and Whitlock 2012). This includes any costs brought on by sexual conflict (Whitlock and Agrawal 2009, Lehtonen et al. 2012, Burke and Bonduriansky 2017). Indeed, both intra- and inter-locus sexual conflict – i.e. sexually antagonistic selection on sex-homologous and sex-heterologous traits, respectively – impose costs to our population’s offspring production (Berger et al. 2016a), which likely drives the cost of sex to be greater than two-fold. Nevertheless, at the very least, our findings indicate that the ability of selection to purge the population’s mutation load is detectable via males, but absent in females, representing a striking difference between the sexes that may partially compensate for the cost of sex.

### Mechanistic understanding

One explanation for why male fitness exhibits greater sensitivity to mutation load is that the fitness consequences of genetic variation in traits under selection are greater (i.e. phenotypic selection is stronger) in males, and/or phenotypic variance in fitness-related traits is more sensitive to mutational input in males (Rowe and Houle 1996). That is, rare partially recessive deleterious mutations may not manifest in female fitness components strongly enough, and/or those female fitness components may not vary enough, to expose those deleterious alleles to selection as readily as in males. Indeed, outbred males exhibited 1.24x the genetic variance in fitness relative to outbred females (Table 1), and males appear to have suffered moderately greater detriments than females from having their partially recessive deleterious alleles revealed by inbreeding/homozygosity (see inbred points above the y=x line in Fig. 1A and Fig. S5). While the present data does not allow us to distinguish between whether male fitness variance is greater because phenotypic selection is stronger in males or because phenotypic traits under selection are more variable in males, these broader characteristics of our population may hold across other animal taxa. Laboratory estimates from insects based on inbreeding depression (Mallet and Chippindale 2011), mutation accumulation (Mallet et al. 2011, 2012, Sharp & Agrawal 2013) and induced mutations (Sharp & Agrawal 2008, Almbro and Simmons 2014, Grieshop et al. 2016) indirectly suggest that males are indeed more sensitive to mutation load than females. Further, meta-analyses show that the opportunity for, and the strength of, selection is generally greater in males than females (e.g. Janicke et al. 2016, Singh and Punzalan 2018).

Quantitative genetic studies of sex-biased genes (genes with sexually dimorphic expression) provide further mechanistic insight to our findings. Male fitness components in *Drosophila melanogaster* are, at least to some extent, determined by the expression levels of genes that typically show male-biased expression (Dean et al. 2018). Further, the expression levels of male-biased genes of *D. serrata* exhibit greater broad-sense heritability than those of female-biased genes (Allen et al. 2018), suggesting that selection could act more efficiently to purge deleterious alleles from any sites that affect the expression levels of male-biased genes. As for how this might affect female fitness, that same study found higher intersexual genetic correlations (*r_MF_*) for expression in male-biased versus female-biased genes (Allen et al. 2018). High *r_MF_* for gene expression or other traits is often interpreted as genetic constraints to sexual dimorphism, possibly imposing sexually antagonistic fitness consequences (Bonduriansky and Chenoweth 2009, van Doorn 2009, Cox and Calsbeek 2009, Connallon et al. 2010, Stewart et al. 2010, Griffin et al. 2013, Ingleby et al. 2014, McGlothlin et al. 2019). However, it is certainly still possible for such male-biased genes to have sexually concordant fitness effects, a core assumption of the “condition dependence” theory for sexually selected traits (Rowe and Houle 1996). Indeed, while mutations in *D. melanogaster*’s male- and female-biased genes had greater detriments to male and female fitness components, respectively, the direction of these effects nevertheless tends to be sexually concordant – i.e. detrimental in both sexes (Connallon and Clark 2011). Of particular relevance to the present study, the large majority of male-biased genes in *C. maculatus* that are expressed in females are actually upregulated in females after mating (Immonen et al. 2017). Thus, much of the mutation load on female reproduction in *C. maculatus* could manifest via the expression of male-biased genes, which the present findings show would be purged more effectively via males. Accordingly, *C. maculatus* male-biased genes show a clear pattern of purifying selection which is not seen in female-biased genes (Sayadi et al. 2019).

While our findings do not offer direct support of “good genes” sexual selection, they do represent evidence of the prerequisite conditions for that process (Zahavi 1975, Lande 1981, Hamilton & Zuk 1982, Grafen 1990, Kirkpatrick and Ryan 1991, Andersson 1994, Kirkpatrick 1996, Kirkpatrick and Barton 1997, Martinossi-Allibert et al. 2019b). Empirical evidence for good genes sexual selection remains scant (Prokop et al. 2012). For good genes sexual selection to work, female choosiness should covary with the “genetic quality” of their mates (i.e. males’ breeding values for fitness; Hunt et al. 2004). Further, that genetic quality should be passed on to both sons and daughters. Lastly, “genic capture” (i.e. polygenic mutation-selection balance; Rowe and Houle 1996, Tomkins et al. 2004) should prevent variation in genetic quality from being depleted. Although our fitness estimates are not a measure of female choice, our findings are consistent with male breeding values for fitness (i.e. genetic quality) reflecting polygenic deleterious mutational variation. While male mating success in this species does seem more to do with male competition than female choice (Savalli and Fox 1999), the kicking behavior that females exhibit may still serve as a baseline level of resistance that enables females to choose the males that are capable of overcoming it (Maklakov and Arnqvist 2009). Further, post-copulatory cryptic female choice (Thornhill 1983, Eberhard 1996, Pitnick et al. 2009, Arnqvist 2014) may comprise a large fraction of male fitness variance in this species (Hotzy et al. 2012, Fritzsche and Arnqvist 2013, Bayram et al. 2019), though there is little evidence for “good genes” effects operating in this context (Bilde et al. 2008, 2009). Thus, whereas only direct selection on female choice has been demonstrated in this system (Maklakov and Arnqvist 2009), the current findings show that the prerequisite genetic architecture is present for indirect “good genes” effects to act in conjunction with direct selection (Kirkpatrick and Barton 1997).

In order for selection via males to yield net benefits to females/populations, thereby contributing to the maintenance of sexual reproduction and female trait preferences via “good genes”, selection in the long run should necessarily act to purge unconditionally deleterious alleles. However, the ability to detect this process in the standing genetic variation or short-term evolutionary outcomes may be overshadowed by genetic variation whose effects are conditional upon sex (Whitlock and Agrawal 2009). Our study population is known to originally harbor sexually antagonistic standing genetic variance in fitness (Berger et al. 2014, 2016a), and as discussed above, the synthetic diallel population analyzed here apparently still does consist of some sexually antagonistic genetic variation (Grieshop and Arnqvist 2018). The present findings are thus a testament to the fact that selection in males against unconditionally deleterious alleles is still detectable and able to promote female offspring production despite the male-imposed detriments of sexually antagonistic genetic variation.

## Acknowledgements

We thank Aneil F. Agrawal (and lab members) and Julia M. Kreiner for comments on earlier drafts of the manuscript, Szymon M. Drobniak and Jacek Radwan for very helpful conversations, Rafael Augusto, Mengjie Fei, and Johanna Liljestrand Rönn for invaluable laboratory technical support, Bo Stenerlöw for access to the irradiation facilities, and Isabelle Glitho for the collection of beetles. This research was supported by the European Research Council (GENCON AdG-294333 to GA), the Swedish Research Council (621-2010-5266 to GA, and 2015-05223 to DB, and 2018-06775 to KG), the NIH (F32AG064883 to PLM), the University of Toronto’s Faculty of Arts and Science (Postdoctoral Fellowship to KG), Stiftelsen för Zoologisk Forskning (to KG), and a Liljewalch’s Resestipendier (to KG).

## Authors’ contributions

KG, DB and GA conceived of and designed the experiment. KG conducted the experiment. KG, DB, and PLM conducted the statistical analyses. KG wrote the first draft of the manuscript. All authors contributed to editing the manuscript.

## Data availability

The raw data, R code and a dependency, which together enable all analyses, procedures and figures to be reproduced, have been submitted for review along with the model output files under the reported settings; these are available at Dryad accession number: XXX.

## Appendix

**Table A1:**
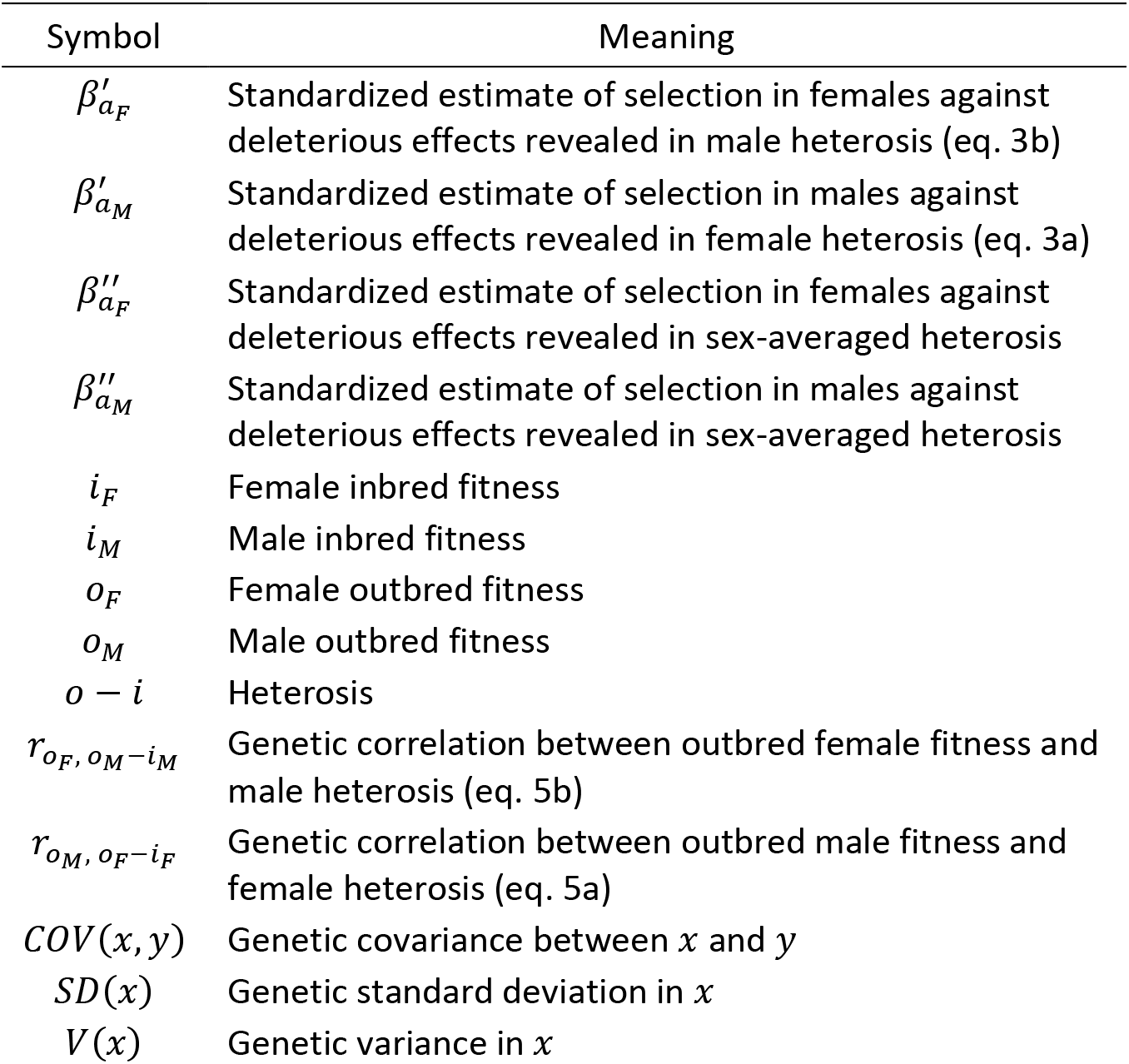
Frequently used symbols from the main text.

## A2. Regarding the role of parental-effects and sex-chromosomes

## A2 Rationale

There is the potential that non-genetic parental effects of each strain could partly confound our additive genetic interpretation of the *o_M_* and *o_F_* estimates. That is, the *o_M_* and *o_F_* estimates of each point on Fig. 1C stem from replicate observations of the same genetic crosses, meaning that, in one extreme limit, differences among them could be attributable to the inheritance/transfer of phenotypic condition from those same recurrent strains to their F_1_ offspring rather than the inheritance of the genetic makeup of those strains, *per se*. Such “condition transfer” via parent-of-origin effects (e.g. epigenetics) may be a common adaptive feature of many organisms (Bonduriansky and Crean 2018). Further, genes with male-biased gene expression, which we discuss as possibly mediating our findings (see *Mechanistic understanding*), may be likewise particularly relevant to condition transfer via parent-of-origin effects, as they have been shown to exhibit elevated condition-dependent expression in *Drosophila melanogaster* (Wyman et al. 2010). In addition to non-genetic parental effects, our main finding of sex differences in the relationship between heterosis and outbred breeding values for fitness could be partly attributable to the mutation load carried only by males on the Y-chromosome, and/or that revealed only in males on their unmasked hemizygous X-chromosome, whereas X-chromosome heterozygosity might mask these effects in females.

We sought to address these potential concerns. Diallel data lend themselves particularly well to identifying these effects via contrasts of reciprocal full siblings – pairs of crosses that are autosomally identical but have inherited their sex-chromosomes, mitochondria, cytoplasm, and other epigenetic information from opposite strains (e.g. the F_1_ offspring of a strain-1 father and strain-2 mother are reciprocal full siblings with the F_1_s of a strain-2 father and strain-1 mother, see Fig. S1B). These parent-of-origin effects were identified via diallel variance partitioning (after Shorter et al. 2019) and removed from the fitness data, yielding a new “Y-adjusted” fitness variable that was subject to the same analyses reported in the *Methods*.

## A2 Methods

To estimate and remove the contribution of strain-specific parental sex effects (*m* and *ϕ*^*m*^) and asymmetric epistatic effects (*w* and *ϕ*^*w*^) from our fitness phenotype, *y*, resulting in *y*^adjusted^, we implemented an updated version of the linear mixed model previously used in Shorter et al. (2019), using the R software package MCMCglmm (Hadfield 2010). We used the BayesDiallel ‘fulls’ (full, sexed, ‘BSabmvw’) model to estimate the contribution of parental strains and their various effects as described in (Lenarcic et al., 2012), where for an individual *i* the phenotype is modeled as:

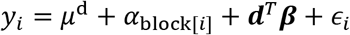

with *ϵ_i_* ~ *N*(0, *σ*^2^). Here, the overall mean and block effects are modeled by *μ*^d^ and *α_block_*, respectively. For dam *j*, sire *k,* and sex *s*, with indicator functions *ψ*, the diallel effects, **d**^*T*^***β***, are estimated by

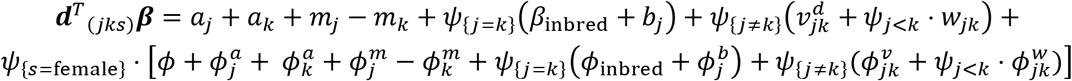

where for all *j* strains, the strain-specific effects are modeled marginally for additive effects as 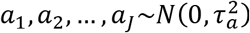, parental sex effects as 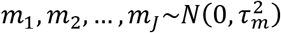, inbred effects 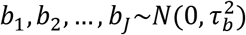 and similarly for sex-specific versions of these effects classes (*ϕ*^*a*^, *ϕ*^*m*^, *ϕ*^*b*^). The strainpair-specific effects for all unique *j*-by-*k* non-inbred combinations, are modeled similarly for symmetric epistatic (*v*^*d*^), asymmetric epistatic (*w*), and sex specific versions (*ϕ*^*v*^ and *ϕ*^*w*^, respectively), e.g. as marginally 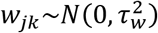 for asymmetric epistatic effects.

The priors for the five fixed effects (*μ*^d^, *a*_block_, *β*_inbred_, *ϕ* (overall female), and *ϕ*_inbred_) are each set to a vague normal distribution, fixed. effect ~ *N*(0,10^3^), and the priors for variance of the residuals (*σ*^2^) and for the variance for each class of effects (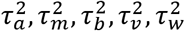 and sex-specific versions), are set as a weakly informative Inverse-Wishart with V=2, and nu=0.002, equivalent to an inverse gamma prior with shape and scale of 0.001. The estimates are based on models that were run for 17,000 iterations after 2,000 iterations of burn-in.

After obtaining stable estimates, e.g. 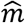 for parental effects, for all strains and strainpairs, the original fitness phenotypes are adjusted using the following relationship for each *j*, *k*, and *s*:

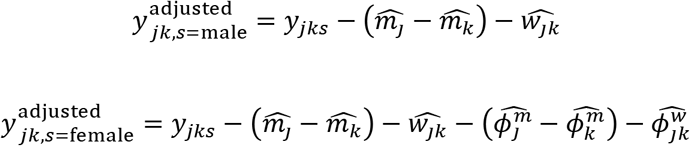

These adjusted Y fitness values are then modeled in the same way as the original fitness values in the main text (see *Methods*).

## A2 Results

The results we obtained using the Y_adjusted fitness values were highly consistent with those reported in the main text. The model fits (see equation 2, *Methods*) before and after removing the parental effects were negligibly different (DIC = −1219.80 and DIC = −1222.275, respectively). The standardized **H** matrix estimated after removing the parental effects from the data (Table A1) was qualitatively similar to before (Table 1). Resampling this model revealed that heterosis in males and females was still highly positively correlated (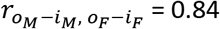, (0.67, 0.96), *P* < 0.001). Likewise, male outbred breeding values were highly negatively correlated with heterosis in females (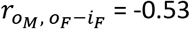, (−0.84, −0.14), *P* = 0.014), whereas female outbred breeding values were not correlated to heterosis in males (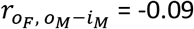, (−0.44, 0.25), *P* = 0.674). Thus, our analysis suggests that non-genetic parental effects, sex-chromosome effects, and their epistatic interactions with autosomal genetic variation do not contribute to our main findings. For any parental effects variance to still be present in the Y-adjusted fitness variable, and hence still clouding our interpretation, the pattern of strain-specific non-genetic inheritance would need to be identical between reciprocal full-sibling contrasts (as well as between the sex-specific, i.e. son-daughter, contrasts of those reciprocal full-sibling differences). We argue that this is highly unlikely considering that parental effect variance manifests in fathers and mothers via fundamentally different and highly asymmetric pathways (e.g. cytoplasmic, mitochondrial, and sex-chromosome inheritance).

**Table A2.** **H** matrix (with CIs and residual variances) estimated for relative Y-adjusted fitness (i.e. relative fitness after removing parental effects; see *Methods* and S2). Genetic variances (and residual variances) on the diagonal, their covariances in the lower triangle, and their Pearson’s correlation coefficients (*r*) in the upper triangle. Covariances and correlations with 95% credibility intervals excluding zero are bolded.

| | $i_F$ | $O_F$ | $i_M$ | $O_M$ |
| --- | --- | --- | --- | --- |
| $i_F$ | 0.0109<br>(0.004, 0.021)<br>(0.0257) | 0.35 | **0.82** | 0.48 |
| $O_F$ | 0.0011 | 0.0015<br>(<0.001, 0.003)<br>(0.0149) | 0.16 | -0.02 |
| $i_M$ | **0.0106** | 0.0002 | 0.0195<br>(0.006, 0.038)<br>(0.0857) | **0.70** |
| $O_M$ | 0.0017 | $-1.8 \times 10^{-5}$ | **0.0030** | 0.0019<br>(<0.001, 0.004)<br>(0.0836) |

## Appendix 3: Regarding sexually antagonistic variants and heterosis

In an extreme case, in principle, the present synthetic diallel population may be largely fixed throughout all 16 inbred strains for female-benefit sexually antagonistic alleles at many genetic loci due to excess lineage extinction among inbred lines that originated from isofemale lines that were enriched for male-benefit/female-detriment sexually antagonistic genetic variants (Grieshop et al. 2017). Yet, some remaining sexually antagonistic polymorphisms demonstrably still segregate among the inbred strains (Grieshop and Arnqvist 2018). Under one scenario, some strains could be completely fixed for the female-benefit alleles across all sexually antagonistic loci, while other strains may be fixed for most, but not all, female benefit alleles, and thus fixed for male-benefit alleles at the remining sites. Under diminishing returns epistasis (Whitlock et al. 1995, Martin et al. 2007, Berger and Postma 2014), which is theoretically predicted among polymorphic sexually antagonistic sites (Arnqvist et al. 2014), strains fixed for female-benefit alleles throughout all sexually antagonistic loci would tend to have relatively low male fitness (*o_M_*) and relatively high measures of male heterosis (*o_M_*-*i_M_*) owing to the recruitment of male-benefit alleles (upon outcrossing with strains that are fixed for some male-benefit alleles) in an otherwise male-detriment genetic background. By contrast, strains that have some male-benefit alleles fixed would have relatively high male fitness and relatively low measures of male heterosis, as they have less to gain than strains that are fixed for female-benefit alleles across all sexually antagonistic loci (Arnqvist et al. 2014). Female fitness and heterosis would be relatively unaffected in this context because they are either completely or mostly saturated for alleles that benefit their fitness, making the recruitment of additional female-benefit alleles in an otherwise female-benefit background relatively ineffectual (Arnqvist et al. 2014). At first glance, this explanation seems to match our results in that fitness would be negatively correlated to heterosis in males but not females. However, our main finding is specifically between male fitness (*o_M_*) and *female* heterosis (*o_F_*-*i_F_*) (Fig. 1C). Further, *o_M_*-*i_M_* ≈ *o_F_*-*i_F_* (Fig. 1B). Neither of those findings are compatible with an explanation based on sexually antagonistic genetic variation (and sexually antagonistic diminishing returns epistasis) underlying heterosis.

## Supporting Information

**Table S1:** Comparison and rationale for using opposite-sex heterosis for point estimates and sex-averaged heterosis for sex-differences. Using opposite-sex heterosis is the most technically correct approach for the point estimates and assessment of whether they are different from zero, but causes male and female estimates to not be directly comparable. Using sex-averaged heterosis renders the sex-specific estimates directly comparable – and is therefore the most technically correct assessment of fold differences and significance between the sexes – but the estimates may be biased due to shared measurement error between fitness and heterosis. Estimates and values used to support the conclusions are bolded.

| Description | Heterosis | Sex | Symbol | Estimate (CIs) | *P* value | Fold difference | Sex-diff. *P* value |
| --- | --- | --- | --- | --- | --- | --- | --- |
| Genetic standardized selection | Opposite-sex | **Male** | $\beta'_{a_M}$ | **-0.0125**<br>**(-0.031, -0.003)** | **0.008** | 8.7x | 0.11 |
| | | **Female** | $\beta'_{a_F}$ | **-0.0014**<br>**(-0.013, 0.009)** | **0.672** | | |
| | Sex-averaged | Male | $\beta''_{a_M}$ | -0.0133<br>(-0.029, -0.004) | 0.008 | **3.7x** | **0.104** |
| | | Female | $\beta''_{a_F}$ | -0.0036<br>(-0.011, 0.007) | 0.558 | | |
| Genetic correlation | Opposite-sex | **Male** | $r_{o_M, o_F - i_F}$ | **-0.59**<br>**(-0.81, -0.11)** | **0.008** | - | 0.132 |
| | | **Female** | $r_{o_F, o_M - i_M}$ | **-0.14**<br>**(-0.40, 0.28)** | **0.672** | | |
|  | Sex-averaged | Male | - | -0.46<br>(-0.79, -0.12) | 0.008 | - | **0.12** |
|  |  | Female | - | -0.06<br>(-0.37, 0.17) | 0.558 |  |  |

**Fig. S1:**
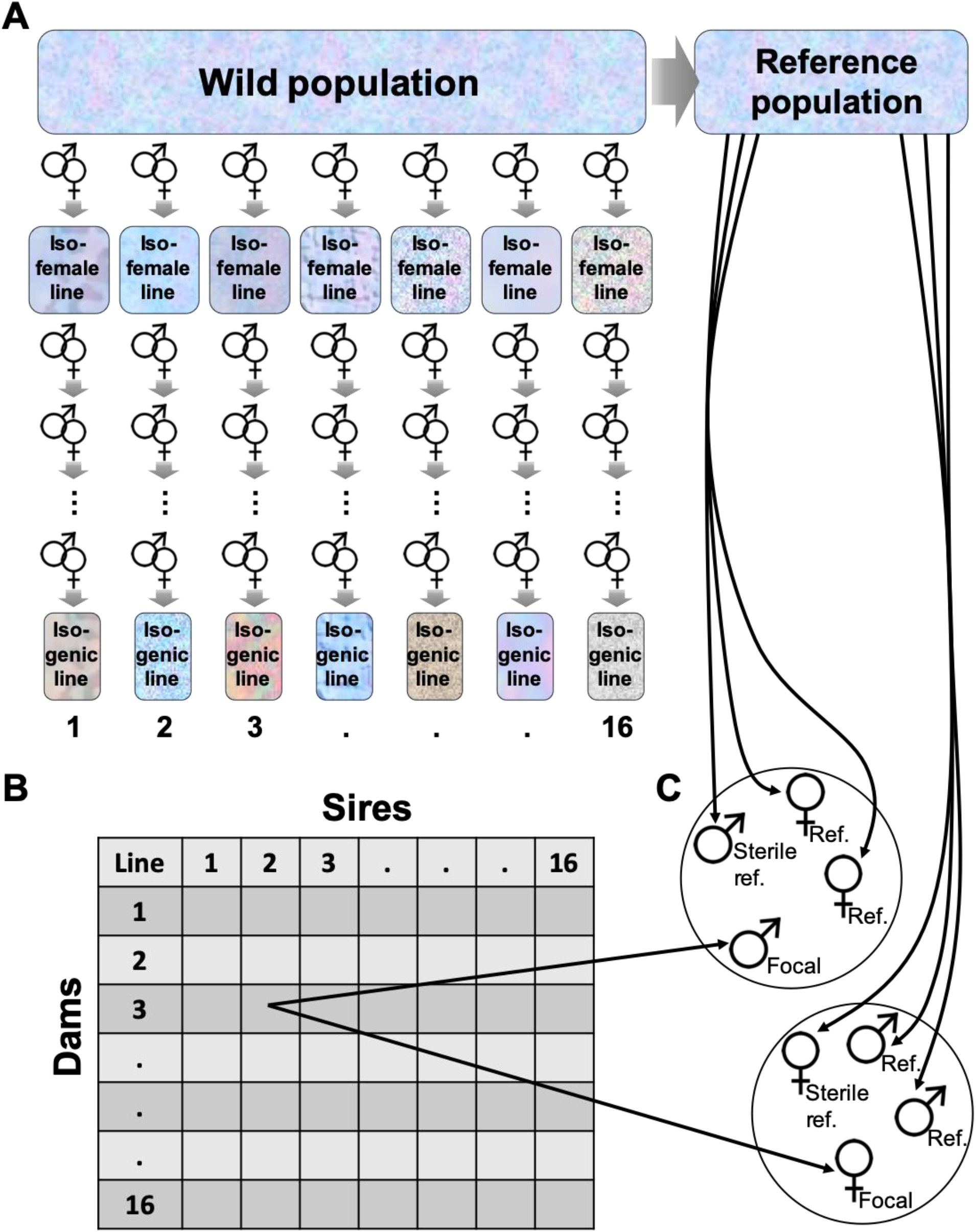
Lome population history and experimental design. (A) A wild population was divided into an outbred laboratory reference population and 41 isofemale lines (Berger et al. 2014). The latter were further inbred (Grieshop et al. 2017) to obtain the 16 inbred/isogenic lines used presently (Grieshop and Arnqvist 2018). Genetic diversity is depicted by color and texture. (B) A full diallel cross (Lynch and Walsh 1998) among the 16 inbred strains, where F_1_ inbred parental selfs (*i*) are on the diagonal and outcrossed F_1_s (*o*) are on the off-diagonal. (C) Replicate F_1_ male and female fitness estimates included a focal F_1_ individual (e.g. outbred (*o*) F_1_s from a strain-2 sire and strain-3 dam), a sterile same-sex competitor from the reference population, two (fertile) opposite-sex competitors from the reference population, and ca. 100 *V. unguiculate* seeds in a 90 mm ø petri dish. These beetles were left to interact/mate/oviposit for the duration of their lifetime, and F_2_ offspring counts = focal individuals’ fitness.

**Fig. S2:**
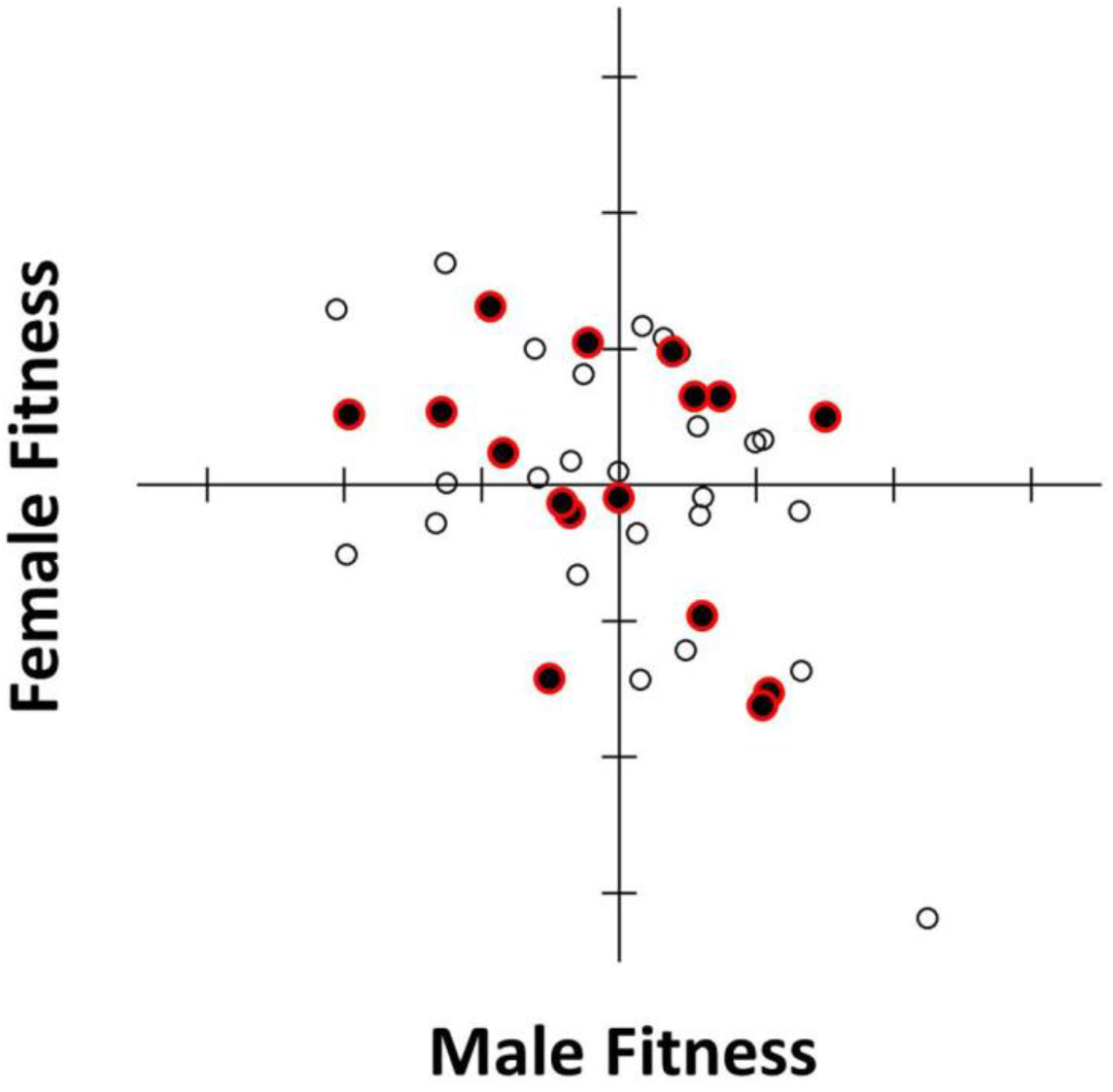
The raw-means *r*_MF_ of the isofemale lines (see Fig. S1; Berger et al. 2014). Those from whence the present 16 inbred strains stem are outlined in red and filled in black. Because all 20 replicate inbreeding lineages stemming from some of the most male-benefit/female-detriment isofemale lines went extinct prior to completing the full inbreeding program (Grieshop et al. 2017), the present 16 inbred strains were chosen from throughout the isofemale line *r*_MF_ with the aim of countering that bias.

**Fig. S3:**
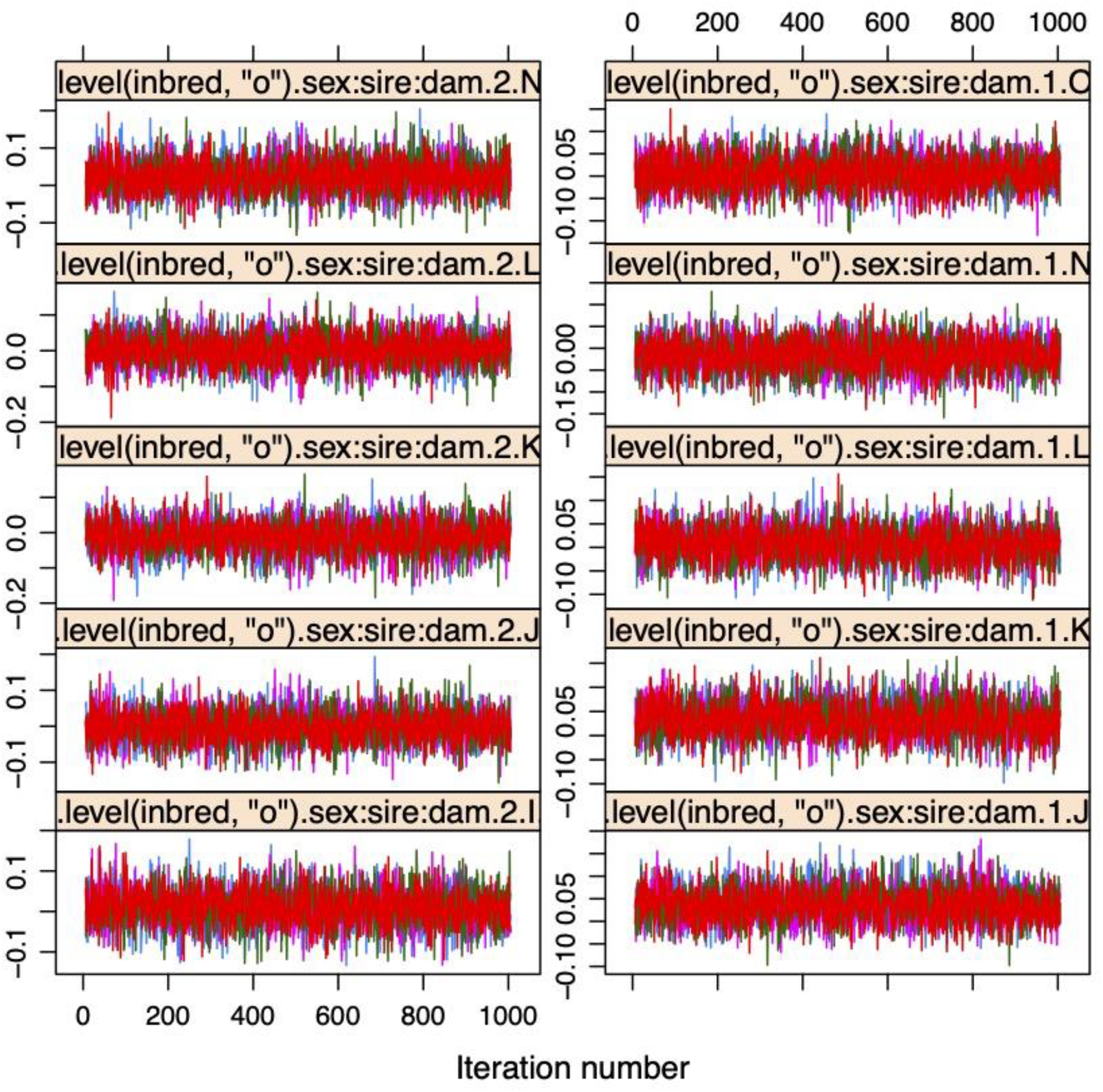
Results the Gelman-Rubin analysis, demonstrating good mixing of four independent MCMC chains. Shown is a representative random sample of ten parameters, out of the total 780 parameters (plus an intercept) that were estimated in our model. Other diagnostic output from this analysis is available in the R script (starting on line 110).

**Fig. S4:**
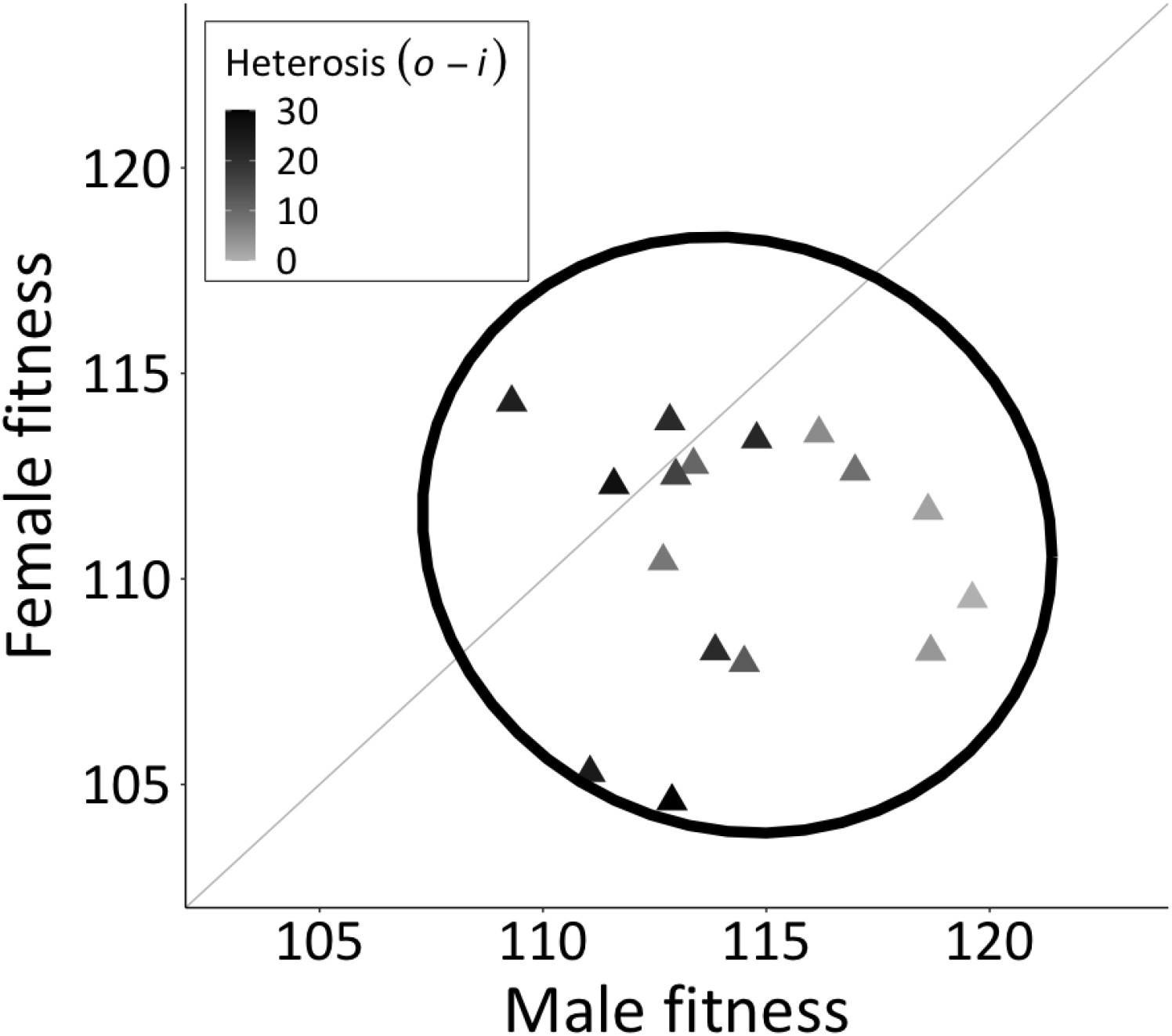
Outbred (*o*) breeding values from Fig. 1A shaded by sex-averaged heterosis. An informal depiction of our main finding is apparent in that heterosis is clearly distributed along a horizontal gradient, according to male breeding values for fitness, but is clearly not distributed along the vertical dimension. Thus, sex-/strain-specific heterosis, the degree to which inbred strains benefit from having their rare partially recessive deleterious mutations covered up by heterozygosity, is reflected in those strains’ outcrossed males, but not their outcrossed females.

**Fig. S5:**
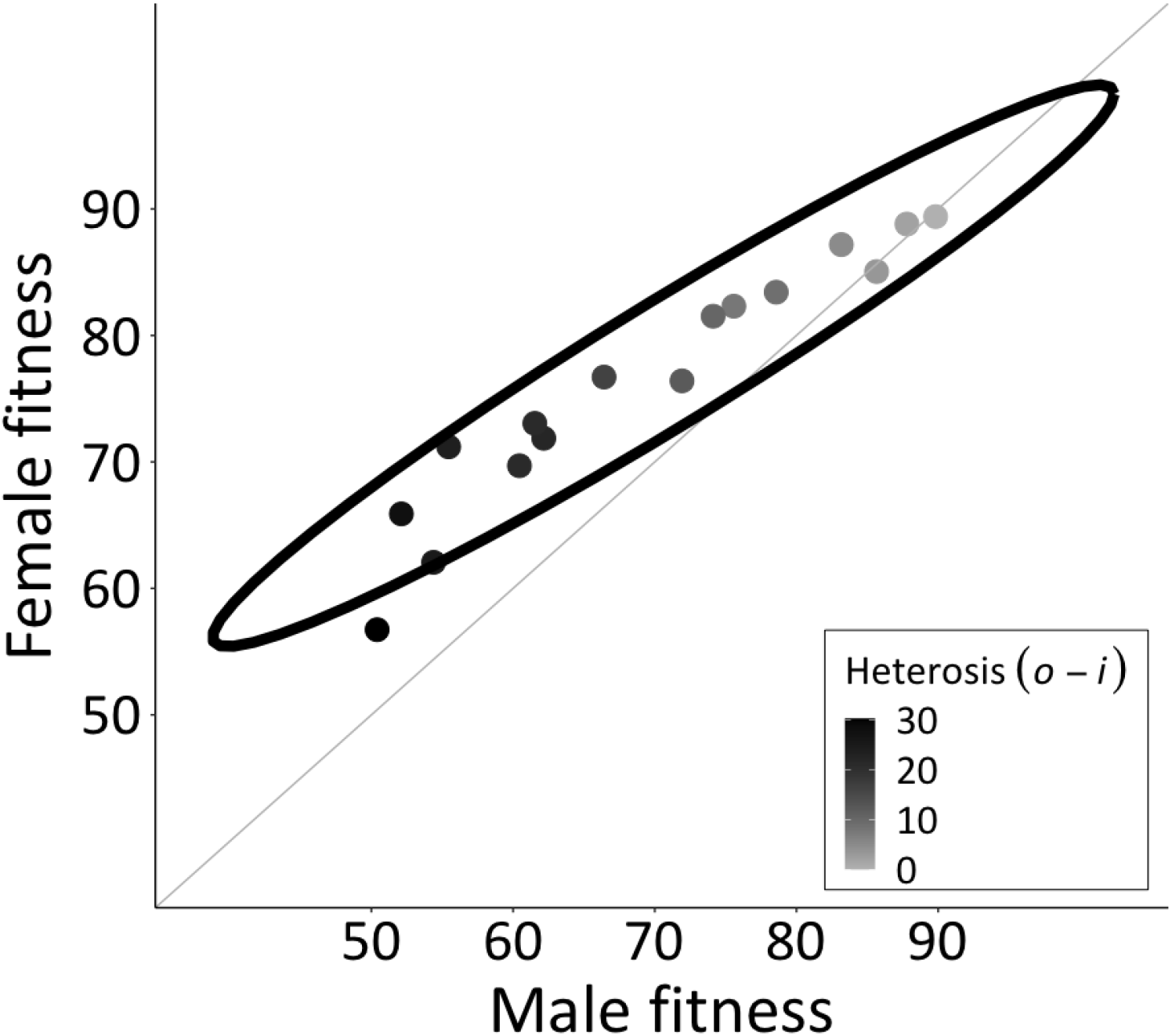
Inbred (*i*) breeding values from Fig. 1A shaded by sex-averaged heterosis. By definition, strains with greater inbred fitness experience less heterosis. Male fitness is more negatively impacted by inbreeding/homozygosity than female fitness, as the large majority of these strains’ inbred breeding values lie above the y=x line.

**Fig. S6:**
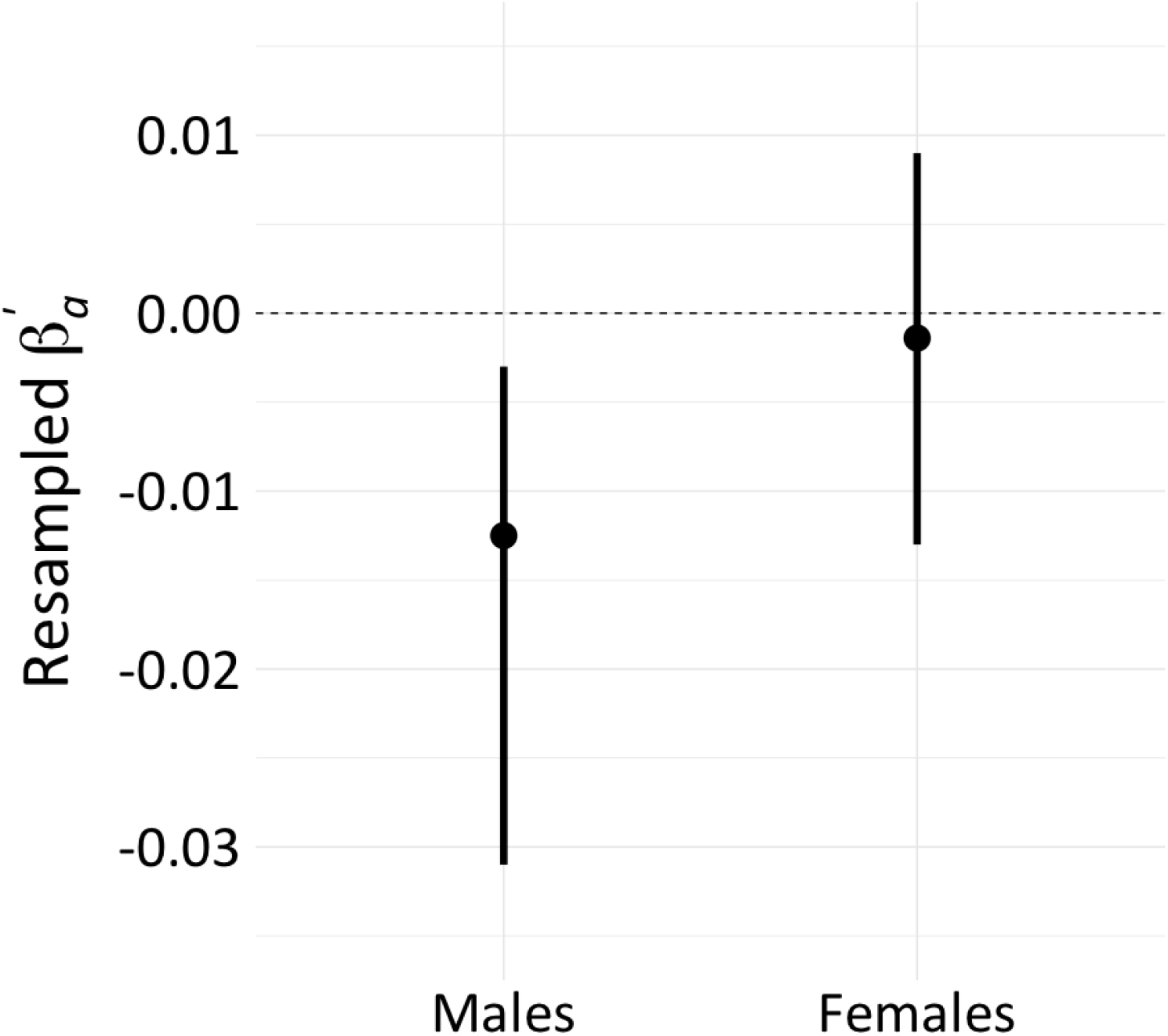
Resampled point estimates and 95% credibility intervals of 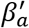 for males 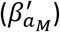 and females 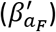.

